# Comparative Analysis of Phytochemical Constituents of *Ginkgo Biloba* Flowers and Leaves, and Evaluation of Their Biological Activities

**DOI:** 10.1101/2025.01.26.634959

**Authors:** Yu-Ying Wang, Rui-Hong Li, Xin Sun, Zi-Ming Xia, Guang-Jie Zhang, Bin Li, Ying Tian, Min Li, Shu-Chen Liu

## Abstract

**Backgrounds:** *Ginkgo biloba* L. has attracted much attention for its unique chemical composition and pharmacological properties, and most of the research focused on *Ginkgo biloba* leaves (GBL). Our preliminary research found that *Ginkgo biloba* flowers (GBF) were superior to GBL in terms of anti-radiation activity, but the underlying cause of this discrepancy in activity remains elusive.

**Objectives:** The aim of this study was to systematically compare the chemical and nutritional composition of *Ginkgo biloba* flowers and leaves, to further elucidate the material basis of the medicinal and nutritional value of *Ginkgo biloba* flowers, and to conduct further comparative studies on the anti-ferroptosis activity and anti-radiation activity of the two.

**Methods:** In this study, the chemical constituents of GBF and GBL were identified by UPLC-Q-orbitrap HRMS, and the contents of amino acids, fatty acids, inorganic elements, purines, and ceramides of both were determined. Meanwhile, the anti-iron death activity and anti-radiation activity of GBF and GBL were comparatively studied by cellular experiments.

**Results:** The results showed that the content of chemical components such as flavonoids and other nutrients such as amino acids, fatty acids, and ceramides were richer in GBF than in GBL, and GBF was indeed superior to GBL in terms of anti-radiation and anti-ferroptosis activity.

**Conclusion:** Through in-depth analyses of the chemical and nutritional compositions of *Ginkgo biloba* flowers and leaves, the present study reveals the possible material basis for the stronger activity of GBF and provides theoretical support for its application in nutraceuticals and pharmaceuticals, especially in the development of functional nutraceuticals and anti-radiation drugs with higher health benefits.

## Introduction

*Ginkgo biloba* L., also called maidenhair tree, is a rare species of Mesozoic relict and the sole species in the class Ginkgopsida, also known as the “living fossil”. In recent years, the chemical composition and pharmacological properties of GBL have been the subject of considerable research(Liu et al. 2022). The GBF belonging to the gymnosperm group, has been observed to produce male globular flowers(Li 2019).

Radiotherapy has been pivotal in cancer therapy since its discovery, but Ionizing radiation in radiotherapy can cause a variety of serious acute and chronic complications. Ionizing radiation is capable of triggering intracellular oxidative stress, leading to a high production of reactive oxygen species (ROS). This state of oxidative stress promotes the onset of ferroptosis because excess ROS induces the degradation of ferritin, releasing Fe, while attacking polyunsaturated fatty acid phospholipids in the cellular membranes, triggering lipid peroxidation, which is one of the key features of ferroptosis. In addition, ionizing radiation-induced DNA damage inhibits the expression of SLC7A11, a key part of cystine/glutamate transporter proteins, sustains GSH depletion and inhibits GPX4, further promoting ferroptosis. Thus, there is a close and complex relationship between ionizing radiation, oxidative stress, and ferroptosis(F. Wang et al. 2024) (Jiao, Cao, and Liu 2022).

It has been found that it is possible to inhibit ferroptosis and oxidative stress to achieve anti-radiation effects(Berry et al. 2024a). Our group found that *Ginkgo biloba* flower has a certain antioxidant effect, and found that the flavonoids and other components in *Ginkgo biloba* flower also have good anti-ferroptosis activity, and a comparative study of the anti-radiation activity of *Ginkgo biloba* flower and leave found that *Ginkgo biloba* flower is better than leave in terms of anti-radiation activity (Li 2019; Li et al. 2021; Li, Zhang, and Zhang 2020). However, the research on the chemical composition and biological activity of GBF is still in the preliminary stage, and the underlying mechanism responsible for this discrepancy in activity between the two remains unclear. The discrepancies in the chemical compositions of the two were not readily distinct, necessitating a comparative analysis of the chemical compositions of GBL and GBF to facilitate the optimal utilization of GBF and to elucidate the broader activity and the material basis of GBF.

In this study, the chemical constituents in GBL and GBF were identified through the use of UHPLC-Q-Orbitrap HRMS. Furthermore, the contents of amino acids, fatty acids, inorganic elements, purines, and ceramides in the GBF and GBL were also determined. We also conducted a comparative study on whether GBF and GBL can exert anti-radiation activity by modulating ferroptosis and whether GBF has a superior effect to GBL in inhibiting ferroptosis. This study will establish a foundation for the comprehensive development and utilization of GBF, enrich the chemical composition and biological activity research of *Ginkgo biloba*, and provide a foundation for the design and development of innovative drugs.

## 1 Materials and Methods

### 1.1 Plant material

*Ginkgo biloba* flower samples were obtained from the Tancheng Ginkgo Cultivation Base in Tancheng, Shandong Province, while leaves were procured from Beijing Tongrentang Runfeng Pharmaceutical Co. Following the identification of Ginkgo male flowers and *Ginkgo biloba* leaves by Professor Li Bin, the Beijing Institute of Radiation Medicine associate researcher, the samples were collected.

### 1.2 Chemicals and Reagents

Amino acid, fatty acid internal, and purine standards were purchased from Sigma. (Merck KGaA, German). Ceramide internal standards werepurchased from Avanti Polar Lipids. Water, methanol, acetonitrile, isopropanol, and formic acid were obtained from Thermo Fisher Scientific. Ammonium acetate was obtained from Sigma-Aldrich. Ethyl Alcohol (EtOH), petroleum ether (PE), CHCl3, EtOAc, and n-BuOH (analysis-grade solvents) were obtained from Sinopharm Chemical Reagent Co., Ltd. (Hushi, Shanghai, China).

Dulbecco’s modified Eagle’s medium (DMEM), cell counting kit-8 (CCK-8), antibiotics, and trypsin were purchased from M&C Gene Technology Ltd. (Beijing, China). Fetal bovine serum (FBS) was obtained from Thermo Fisher Scientific (Gibco, USA). Erastin and Ferrostatin-1 (Fer-1) were purchased by Selleck Chemistry (USA). Horse serum was purchased from Kang Yuan Biology (Tian Jin, China).

### 1.3 Cell Model

Highly differentiated rat pheochromocytoma cells (PC12) were purchased from the Chinese Academy of Medical Sciences and cultured in DMEM supplemented with 10% FBS, 5% heat-inactivated horse serum (Kang Yuan Biology, China), and 1% penicillin-streptomycin in a 37°C incubator in a humidified atmosphere with 5% CO_2_.

### 1.4 Determination of chemical composition

#### 1.4.1 Qualitative analysis of total chemical composition

A Vanquish Flex UHPLC chromatograph (Thermo Fisher Scientific, Inc., Waltham, MA, USA) equipped with an ACQUITY UPLC HSS T3 column was used for separation. The mobile phase consisted of water (0.1% formic acid, phase A) and acetonitrile (phase B) with a flow rate of 0.3 mL/min. The column temperature was 40 °C. The following mobile phase gradient was used: 0–1 min: 2% B; 1–14 min: 2–30% B; 14–25 min: 30-100% B; and 25-28 min: 100% B.

The MS data was collected by a hybrid quadrupole orbitrap mass spectrometer (Q Exactive, Thermo Fisher Scientific, Inc., Waltham, MA, USA) equipped with a HESI-II spray probe. The parameters were set as follows: positive ion source voltage of 3.7 kV and negative ion source voltage of 3.5 kV, heated capillary temperature of 320°C, sheath gas pressure of 30 psi, and auxiliary gas pressure of 10 psi.

#### 1.4.2 Purine determination

The Waters ACQUITY UPLC I-CLASS chromatograph equipped with a Waters UPLC HSS T3 (1.8 μm, 2.1 mm×100 mm, column was used for separation. The mobile phases were composed of phase A (water, 0.1% formic acid, 5 mM ammonium formate) and phase B (methanol). The flow rate was 0.3 mL/min, with an injection volume of 5.0 µL and a column temperature of 50 °C. The following mobile phase gradient was used: 0-1min: 1% B; 1-2min: 1-10% B; 2-7min: 10-100% B; 7-7.1min: 100-1% B; and 7.1-9min: 1% B.

The MS data was collected by a Waters XEVO TQ-XS equipped with a HESI-II spray probe. The positive ion source voltage was set at 3.0 kV, while the cone-well voltage was fixed at 10 V. The desolvation temperature was maintained at 500 °C, and the desolvation gas flow rate was set at 1000 L/h.

#### 1.4.3 Ceramide determination

The chromatography system same as 1.4.2. The mobile phases were phase A (acetonitrile: water = 6:4, 0.1% formic acid, 5 mM ammonium acetate) and phase B (isopropanol: acetonitrile = 9:1, 0.1% formic acid, 5 mM ammonium acetate). The flow rate was 0.26 mL/min. The following mobile phase gradient was used: 0–2 min: 0-30% B; 2-12 min: 30-60% B; 12-14min: 60-95% B; 14-18min: 95-100%B; 18-18.1min: 100-0%B; and 18.1-20min: 0%B.

The MS spectrometry system as 1.4.2. The acquisition mode was positive ion (ESI+) with an ion source voltage of 3.0 kV and a temperature of 150 °C; a desolvation temperature of 500 °C and a desolvation gas flow rate of 1000 L/h.

### 1.5 Nutrient content determination

#### 1.5.1 Amino acid determination

The chromatography system same as 1.4.2. The phase comprises two phases: phase A (water, 0.1% formic acid) and B (acetonitrile). The flow rate is 0.5 mL/min. The column temperature was 50°C. The mobile phase gradient was used: 0–0.5 min: 4% B; 0.5–2.5 min: 4-10% B; 2.5-5 min: 10-28% B; 5-6min: 28-95%B; 6-7min: 95%B; 7-7.1min: 95-4%B; and 7.1-9min: 4%B.

The MS spectrometry system is 1.4.2. The positive ion source voltage was set at 1.5 kV, while the cone hole voltage was fixed at 20 V. The desolvation temperature was maintained at 600 °C, and the desolvation gas flow rate was set at 1000 L/h.

#### 1.5.2 Free Fatty Acid Determination

The chromatography system same as 1.4.2. The mobile phases were phase A (with 0.1% acetic acid, 1 mM ammonium acetate) and phase B (isopropanol and acetonitrile (1:1)). The flow rate was 0.26 mL/min, with a sample volume of 5.0 µL and a column temperature of 55°C. The mobile phase gradient was used: 0–2 min: 10-35% B; 2-4 min: 35-85% B; 4-6min: 85-100% B; 6-7.5min: 100%B; 7.5-7.6min: 100-10%B and 7.6-9min: 10%B.

The MS spectrometry system is 1.4.2. The acquisition mode was negative ion (ESI-) with the parameters: ion source voltage-2 kV and temperature 150 °C; desolvation temperature 500 °C and desolvation gas flow rate 1000 L/h; and cone bore voltage 10.0 V and gas flow rate 150 L/h.

#### 1.5.3 Inorganic Element Determination

A mixed standard solution of Inductively coupled plasma-Mass Spectrometry (ICP-MS) was prepared, in which K, Na, Ca, Mg, and Fe were 1000 mg/L, Cu, Zn, Mn, and Al were 10 mg/L, Sr was 100 mg/L. To prepare the internal standard element stock solution (1000 mg/L) Rhodium was diluted to 10 μg/L with 5% nitric acid solution as internal standard solution.

### 1.6 Cell Activity Assay

#### 1.6.1 Anti-Radiation Assay

X-rays were employed with an irradiation dose of 8 Gy and a dose rate of 1.325 Gy/min (voltage 220 kV, current 25 mA). The experiment was divided into six experimental groups: Control, IR, Fer-1, Exrad, GBF, and GBL (All groups except the Control were irradiated).

#### 1.6.2 Anti-Ferroptosis Assay

The experiment was divided into five experimental groups: Control, Erastin, Fer-1, GBF, and GBL (All groups except the Control were treated with 5 μm concentration of erastin). After 24 hours, 10 µL of CCK-8 solution was added to each well, and the incubation was continued for a further 24 hours. The absorbance at 450 nm was then detected using an enzyme marker.

##### 1.6.2.1 Calcein-AM/propidium iodide (PI) staining

PC12 cells were planted in 24-well plates after the placement of cell slides at the bottom of each well. Calcein-AM/propidium iodide (PI) staining (Beyotime, C2015S) was used to distinguish between living and dead cells after the final treatment. After removing the culture medium, 250 µL Calcein-AM/PI working solution was introduced to wells and stained at 37 °C for at least 20 min. Afterward, the solution was discarded, and cells were rinsed softly with PBS once. Finally, the observation and photographing of the cells were conducted through the OLYMPUS CKX53 fluorescence microscope (490nm and 545nm).

##### 1.6.2.2 Detection of intracellular ROS, lipid peroxides (LPO), and Fe^2+^

PC12 cells were seeded at a density of 1 × 10^4^ cells in confocal dishes and pre-protected for 24 h using the complete medium. This was followed by a 6 h stimulation with 5 μM erastin. The cells were then co-incubated at 37 °C for 30 min with DCFH-DA (10 μM), BODIPY™ 581/591 C11 (10 μM), and FeRhoNox^TM^-1 (5 μM) respectively. After triple washing with PBS, finally, the fluorescence signal of DCFH-DA and BODIPY™ 581/591 C11 was detected with the FITC channel, while the FeRhoNox^TM^-1 was assayed with the PE channels. The data was analyzed using the software FlowJo 10.6.2.

##### 1.6.2.3 Determination of intracellular Superoxide dismutase (SOD)

In a 37 °C, 5% CO_2_ incubator, PC12 cells were pretreated with a 6 h erastin (5 μM) stimulation. Cells from each group were then harvested via centrifugation, and total cellular protein was extracted using cell lysis buffer. SOD levels were quantified using their respective cell count normalization kit (Nanjing jiancheng Bioengineering Institute, A001-3-2).

### 1.7 Statistical analyses

Each experiment was independently repeated three times. The data in the bar graphs were manifested as mean ± standard deviation (SD). Statistical differences among multiple groups were conducted by analysis of variance (ANOVA) (\**p* < 0.05, \*\**p* < 0.01, \*\*\**p* <0.001).

## 2 Results and discussion

### 2.1 Determination of chemical composition

#### 2.1.1 Qualitative analysis of total chemical composition

The control database (TCM Pro 2.0, Beijing Hexin Technology Co., Ltd.) and the theoretical database (constructed from literature, public databases, etc.) were searched. The total ion chromatogram (TIC) for GBF and GBL can be seen in **Figure 1**. A total of 123 compounds were eventually identified in GBF and 116 compounds from GBL (**Table 1**). Of these, 73 flavonoids, 12 terpenoids, 10 phenolic acids, 6 alkaloids, and 5 phenylpropanoids were identified in GBF. Other chemical groups were presented as anthraquinone, fatty acids, amino acids, purines, and alkaloids. A total of 65 flavonoids, 10 terpenoids, 8 phenolic acids, 6 alkaloids, and 4 phenylpropanoids were identified in GBL. The content of 57 flavonoids such as ginkgetin, isoginkgetin, and luteolin, and terpenoids such as andrographolide and ginkgolide B was higher than the GBL in GBF, and the content of bilobalide and ginkgolides A, J, and C terpene lactones were lower than the GBL. Among them, bergenin, andrographolide, tiliacein, calycosin, galangin, daidzein, and roburic acid, which were identified in the GBF, but not identified in the GBL, whereas baicalein, nevadensin, luteolin 3’-glucoside, wogonoside, aurantio-obtusin, and casticin were not found in the GBF but were present in the GBL.

**Figure 1.**
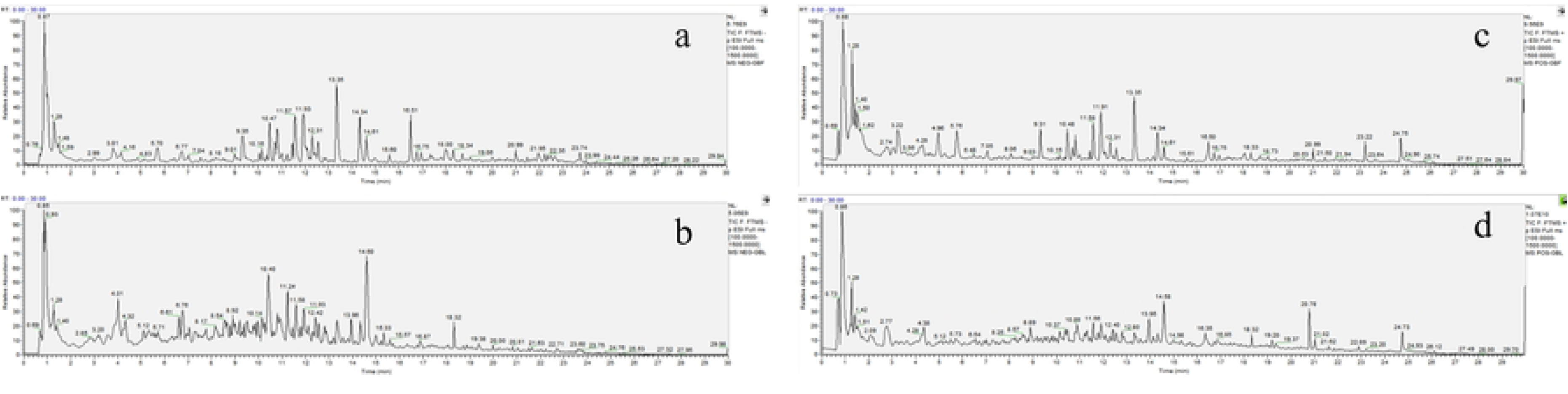
TIC of GBF detected in negative (a) and positive (b) mode. TIC of GBL detected in negative (c) and positive (d) mode.

**Table 1.**
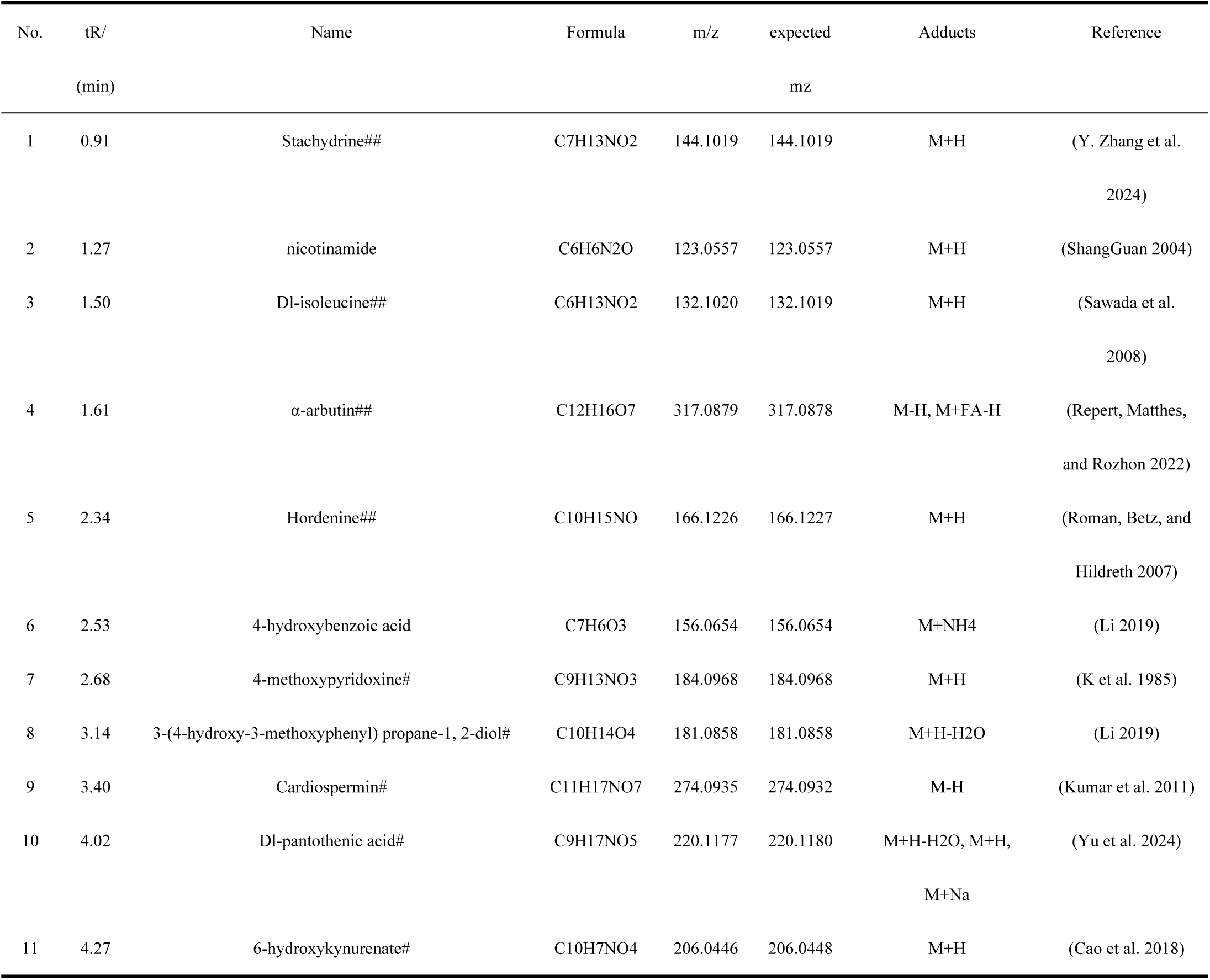
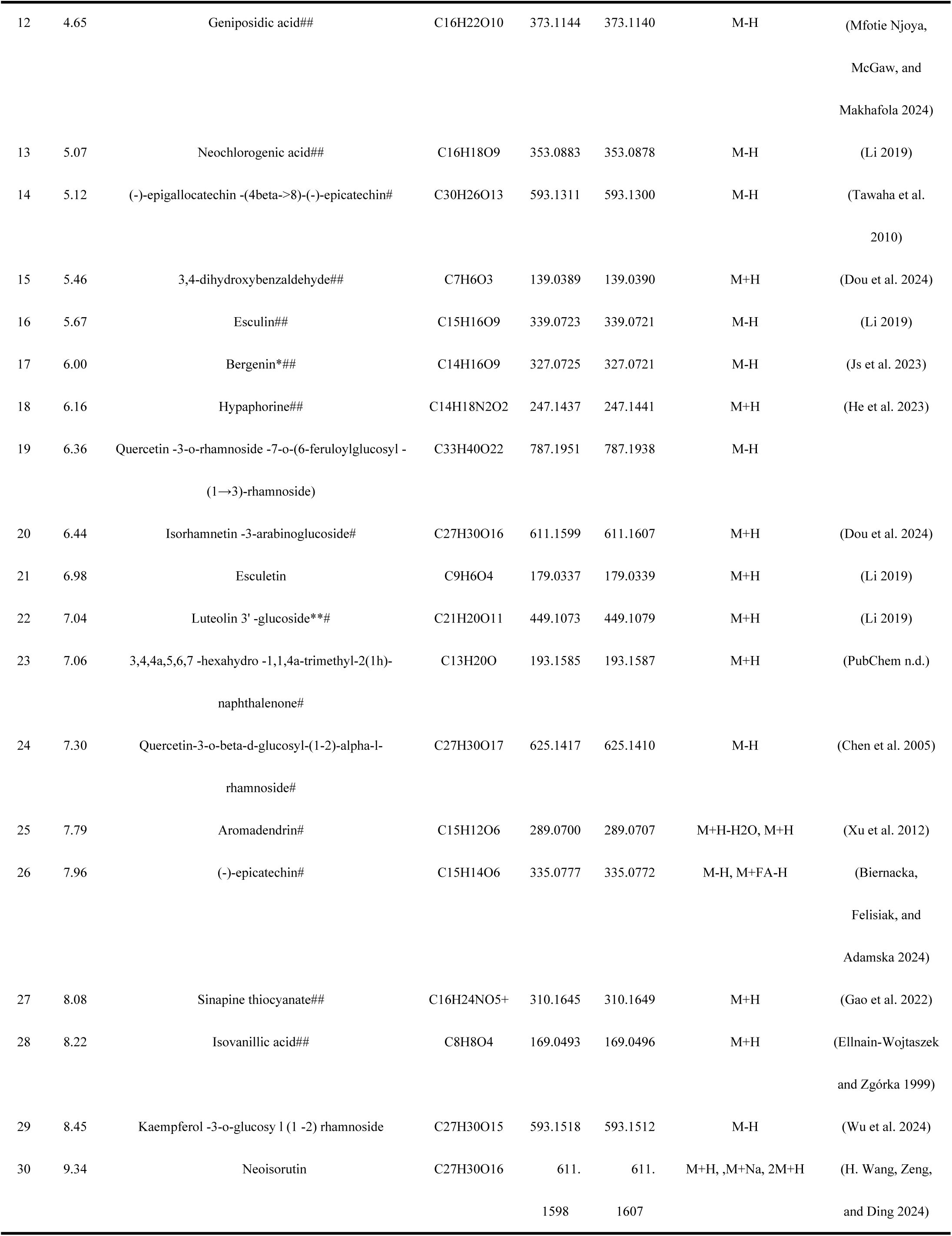
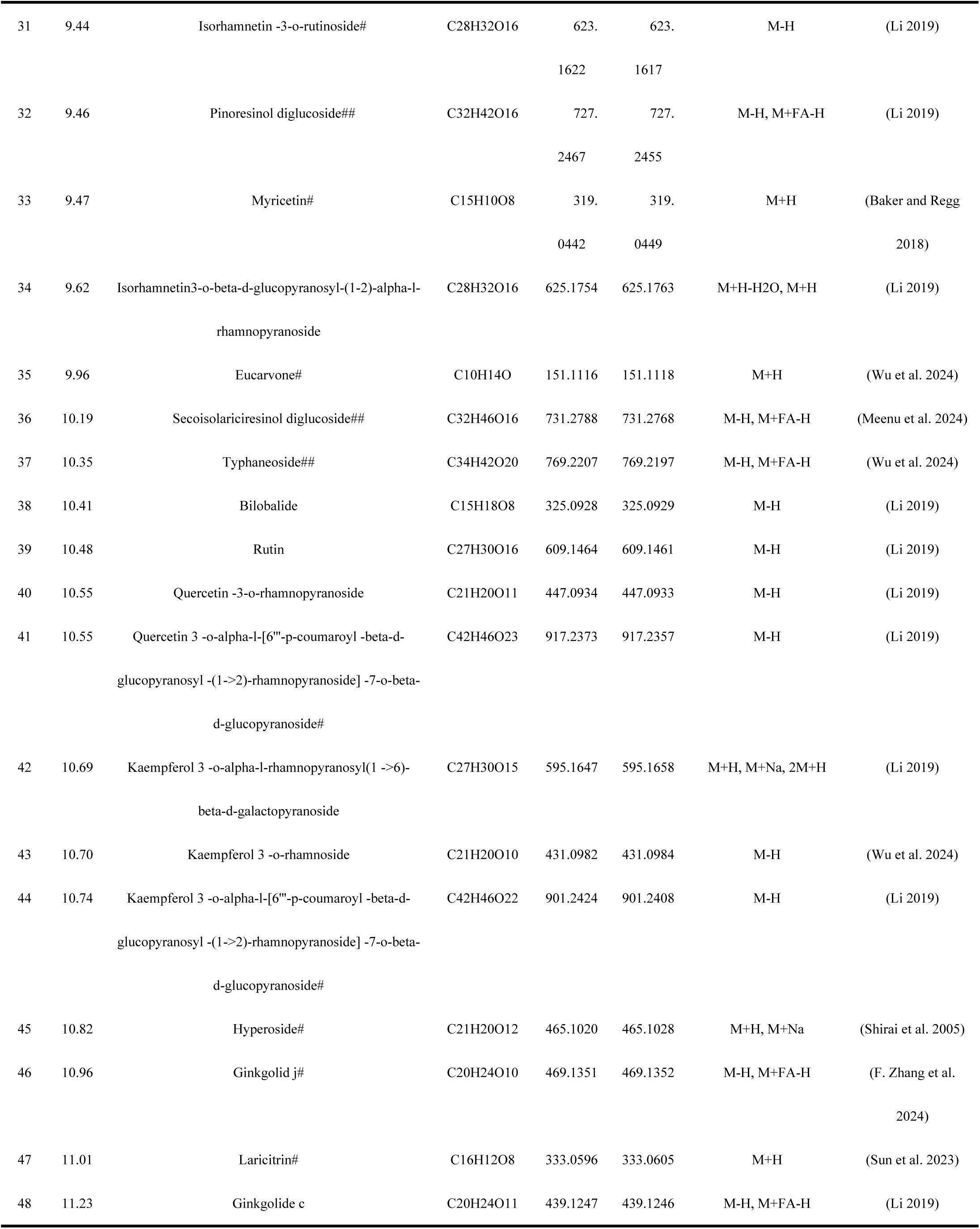
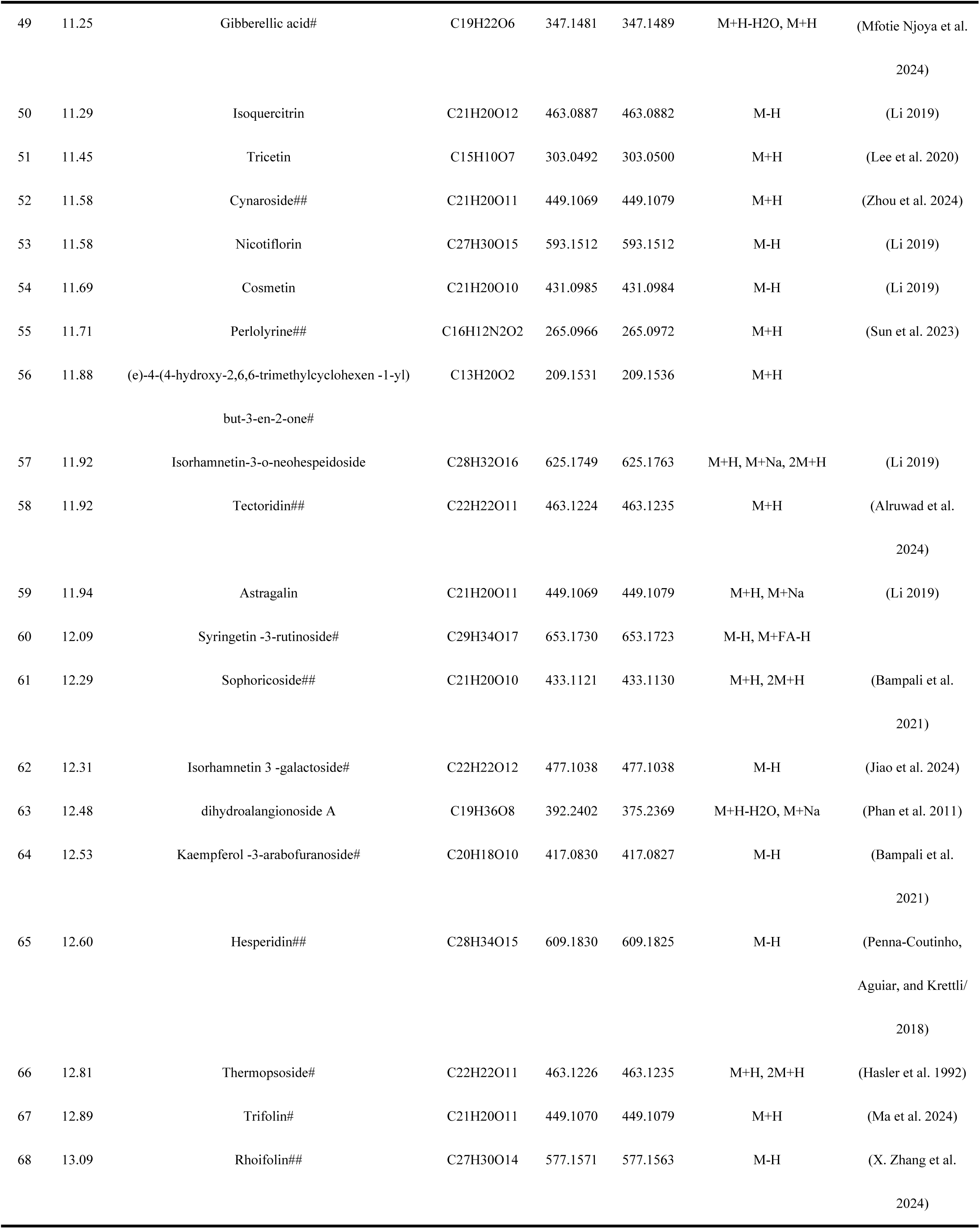
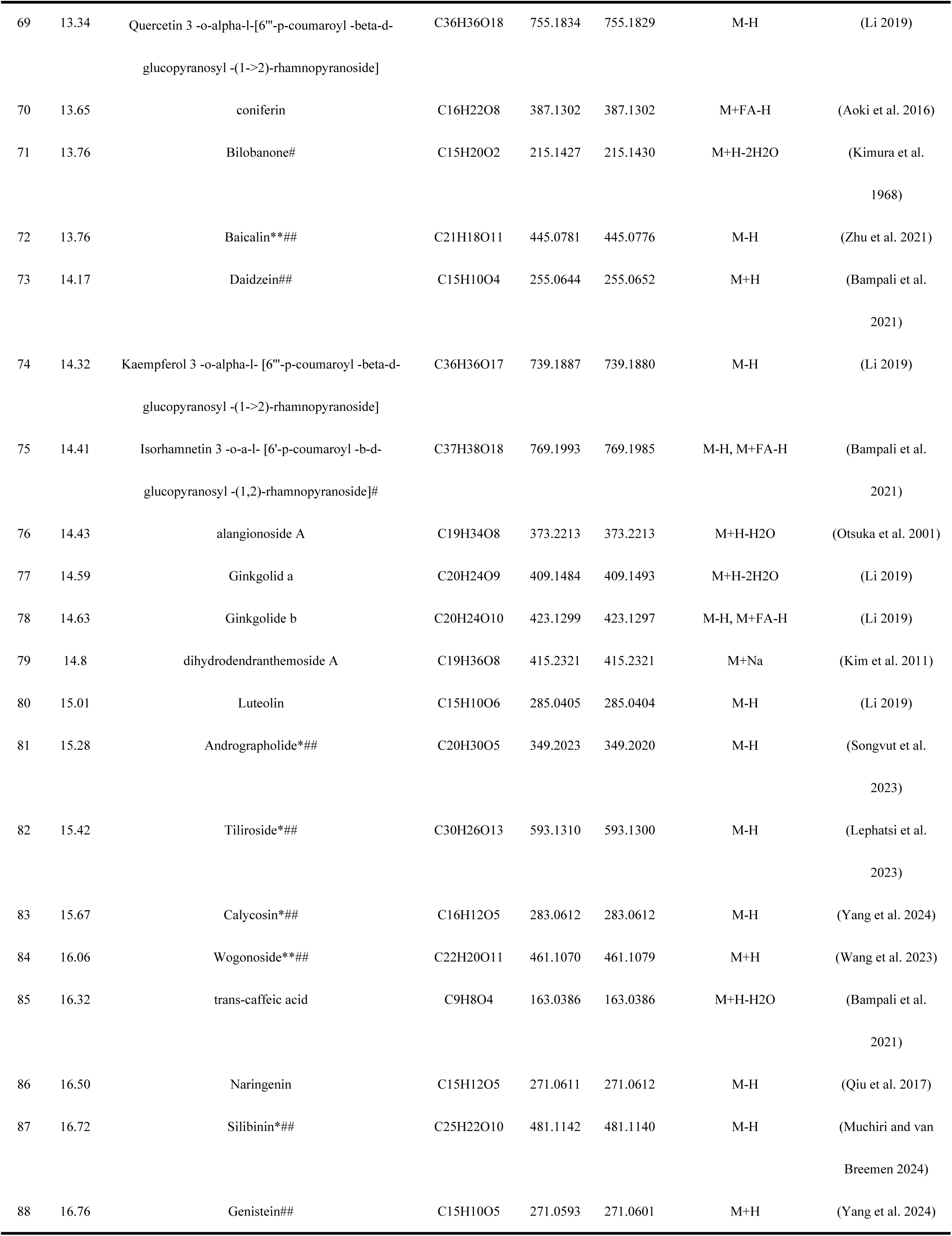
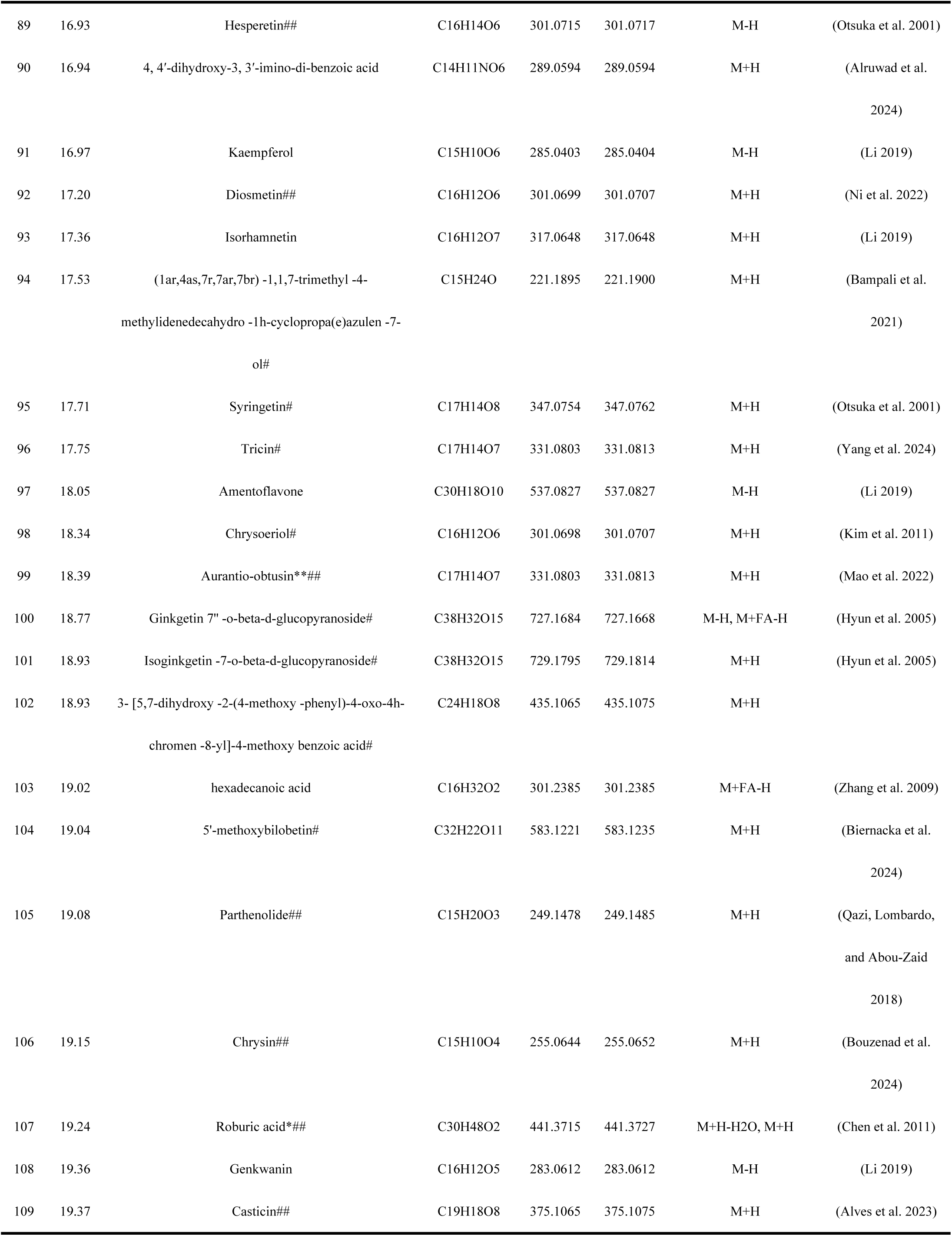
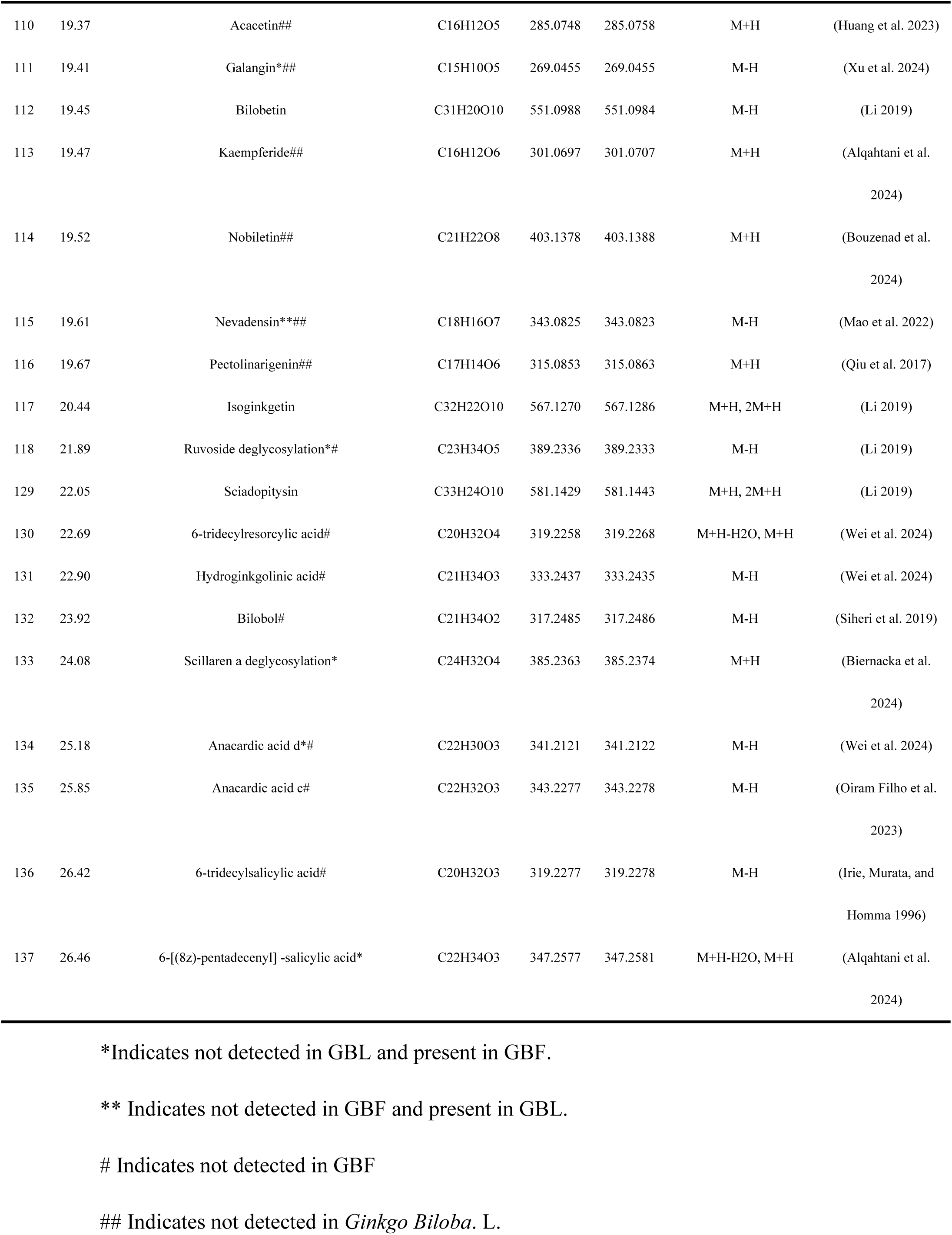
Analysis and identification of chemical constituents in GBF and GBL.

Of these compounds, 41 were identified for the first time in *Ginkgo biloba* (**Table 1**). Furthermore, a total of 76 compounds were identified for the first time in GBF, including 42 flavonoids; 10 terpenoids; 10 phenolic acids; 4 phenylpropanoids; and 6 alkaloids.

At present, the chemical composition of GBL has been identified with a high degree of precision, whereas the chemical composition of GBF remains to be elucidated. In this paper, the preliminary chemical composition identification of GBF and GBL was carried out, and it was found that most of the flavonoids and some of the terpene lactone compounds contained in GBF were more than GBL. Flavonoids appear to be effective in reducing levels of oxidative stress and inflammation in nerve cells and may protect against neurodegenerative diseases (Frandsen & Narayanasamy, 2018). Therefore, it is assumed that GBF has better medicinal value than GBL to some extent.

#### 2.1.2 Purine content determination

Determination of Adenine; Hypoxanthine; Guanine; Xanthine; Adenosine; Inosine; and Guanosine 7 purines were determined (**Table 2**). It can be seen that GBF is also very rich in purines, with total purines in GBF being about 0.8 mg/g, about 10.3 times that of GBL. All purines except hypoxanthine and adenosine were higher in GBF than GBL. Guanosine content reached a maximum of 0.52 mg/g, about 56 times that of GBL. The guanosine was shown to be protective in several experimental models of central nervous system diseases *in vitro and vivo* including ischemic stroke, Alzheimer’s disease, and Parkinson’s disease (Bettio, Gil-Mohapel, and Rodrigues 2016).

**Table 2.** Purines in GBF and GBL in various classes and contents.

|  | GBL | GBF |
| --- | --- | --- |
| Adenine | 24.15±0.43 | 214.77±6.12 |
| Hypoxanthine | 1.16±0.02 | 0.82±0.02 |
| Guanine | 1.23±0.04 | 2.82±0.11 |
| Xanthine | 16.05±0.52 | 43.39±1.16 |
| Adenosine | 23.28±0.43 | 5.71±0.15 |
| Inosine | 2.73±0.09 | 9.97±0.32 |
| Guanosine | 9.34±0.27 | 523.48±24.79 |

#### 2.1.3 Ceramide content determination

A total of 28 Ceramides and 12 Hexosylceramides were determined, and except for 4 Ceramides and 5 Hexosylceramides which were higher in GBL than GBF, the remaining 24 Ceramides and 7 Hexosylceramides were higher in GBF than GBL (**Table 3**). The total ceramide content of GBF was about 12.2 μg/g, which was 6.2 times higher than that of GBL. The cer(d18:2/16:0) content of GBF was about 236 times higher than that of GBL. Studies have shown that ceramide synthesis is required to maintain axonal growth in hippocampal neurons and dendritic growth in cerebellar Purkinje cells. Ceramide also plays different roles at different stages of neuronal development(Schwarz and Futerman 1997).

**Table 3.** Ceramides in GBF and GBL in various classes and contents.

| No. |  | GBL | GBF | No. |  | GBL | GBF |
| --- | --- | --- | --- | --- | --- | --- | --- |
| 1 | Cer (d18:1/14:0) | 0.0225±0.0016 | 0.0230±0.0007 | 21 | Cer (d18:2/24:0) | 0.0059±0.0013 | 0.2771±0.0039 |
| 2 | Cer (d18:1/16:0) | 0.0809±0.0202 | 0.2110±0.0065 | 22 | Cer (d18:2/24:1) | 0.1567±0.0051 | 2.3030±0.0052 |
| 3 | Cer (d18:1/18:0) | 0.2765±0.0081 | 0.0496±0.0035 | 23 | Cer (d18:2/24:2) | 0.0013±0.0004 | 0.0042±0.0003 |
| 4 | Cer (d18:1/19:0) | 0.1497±0.0081 | 0.1720±0.0032 | 24 | Cer (d18:0/16:0) | 0.0161±0.0010 | 0.1732±0.0033 |
| 5 | Cer (d18:1/20:0) | 0.0241±0.0034 | 0.0201±0.0011 | 25 | Cer (d18:0/18:0) | 0.0107±0.0005 | 0.0099±0.0007 |
| 6 | Cer (d18:1/21:0) | 0.0141±0.0010 | 0.0158±0.0003 | 26 | Cer (d18:0/22:0) | 0.0046±0.0004 | 0.0120±0.0001 |
| 7 | Cer (d18:1/22:0) | 0.0296±0.0030 | 0.0331±0.0015 | 27 | Cer (d18:0/23:0) | 0.0036±0.0006 | 0.0055±0.0004 |
| 8 | Cer (d18:1/23:1) | 0.0126±0.0001 | 0.0174±0.0009 | 28 | Cer (d18:0/24:1) | 0.0301±0.0026 | 0.0295±0.0013 |
| 9 | Cer (d18:1/23:0) | 0.0749±0.0049 | 0.0878±0.0033 | 29 | HexCer (d18:1/16:0) | 0.0061±0.0005 | 0.0368±0.0033 |
| 10 | Cer (d18:1/24:1) | 0.0933±0.0086 | 0.1578±0.0059 | 30 | HexCer (d18:1/18:0) | 0.0219±0.0013 | 0.0046±0.0009 |
| 11 | Cer (d18:1/24:0) | 0.1188±0.0132 | 0.1533±0.0009 | 31 | HexCer (d18:1/22:0) | 0.0386±0.0020 | 0.0067±0.0024 |
| 12 | Cer (d18:1/26:1) | 0.0600±0.0080 | 0.0730±0.0051 | 32 | HexCer (d18:1/24:1) | 0.1395±0.0063 | 0.0445±0.0031 |
| 13 | Cer (d18:1/26:0) | 0.0500±0.0072 | 0.0539±0.0017 | 33 | HexCer (d18:1/24:0) | 0.0726±0.0034 | 0.0108±0.0004 |
| 14 | Cer (d18:2/16:0) | 0.0210±0.0009 | 4.9620±0.1339 | 34 | HexCer (d18:1/26:1) | 0.0136±0.0003 | 0.0124±0.0001 |
| 15 | Cer (d18:2/18:0) | 0.0655±0.0020 | 0.2962±0.0094 | 35 | HexCer (d18:2/16:0) | 0.1033±0.0030 | 1.6711±0.0567 |
| 16 | Cer (d18:2/20:0) | 0.0026±0.0011 | 0.1501±0.0036 | 36 | HexCer (d18:2/18:0) | 0.0011±0.0001 | 0.0184±0.0030 |
| 17 | Cer (d18:2/21:0) | 0.0008±0.0005 | 0.0220±0.0006 | 37 | HexCer (d18:2/20:0) | 0.0081±0.0013 | 0.0192±0.0023 |
| 18 | Cer (d18:2/22:0) | 0.0057±0.0008 | 0.2074±0.0113 | 38 | HexCer (d18:2/22:0) | 0.0413±0.0013 | 0.0768±0.0033 |
| 19 | Cer (d18:2/23:0) | 0.0022±0.0008 | 0.0893±0.0029 | 39 | HexCer (d18:2/23:0) | 0.0194±0.0010 | 0.1599±0.0062 |
| 20 | Cer (d18:2/23:1) | 0.0190±0.0006 | 0.2178±0.0103 | 40 | HexCer (d18:2/24:0) | 0.1465±0.0067 | 0.2558±0.0104 |

### 2.2 Nutrient content determination

#### 2.2.1 Amino acid content determination

A total of 38 amino acids were determined (**Table 4**), of which Methionine (MET), CYSTATHIONINE, D-(P) (CYST), Anserine (Ans), Carnosine (CAR), 1-MEHIS, and 3-MEHIS were not detected, and all 31 amino acids were found in GBF except for O-Phosphorylethanolamine (PHETH), the content of GBF was higher than GBL for the remaining 31 amino acids. The GBF amino acid content is as high as 86.9 mg/g, which is about 15.7 times higher than that of GBL, among which the asparagine (ASN) content is the highest at 51.28 mg/g; essential amino acids (EAA) (histidine(HIS), isoleucine(ILE), leucine(LEU), lysine(LYS), phenylalanine(PHE), threonine(THR), tryptophan(TRY), valine(VAL)) content is 10.83mg/g in GBF is 10.83 mg/g, which accounted for 14.28% of the total amino acids, while the EAA content in GBL was only 0.83 mg/g.

**Table 4.** Amino acids in GBF and GBL in various classes and content.

| No. |  | GBL | GBF | No. |  | GBL | GBF |
| --- | --- | --- | --- | --- | --- | --- | --- |
| 1 | HIS | 47.64±3.28 | 1116.37±35.42 | 17 | ALA | 586.15±19.96 | 2858.96±54.48 |
| 2 | OHPRO | 9.19±0.37 | 151.69±6.03 | 18 | γ-GABA | 546.57±23.45 | 3597.12±99.55 |
| 3 | ARG | 111.76±1.53 | 3608.84±136.66 | 19 | OH-LYS | 11.68±0.38 | 51.65±2.29 |
| 4 | ASN | 233.74±8.02 | 51287.55±1623.03 | 20 | AMADP | 11.91±0.4 | 51.45±0.48 |
| 5 | PHETH | 318.55±9.28 | 144.36±6.02 | 21 | PRO | 2085.55±57.97 | 4091.48±81.63 |
| 6 | TAU | 4.09±0.23 | 10.17±0.87 | 22 | β-GABA | 14.36±1.32 | 22.06±1.6 |
| 7 | GLN | 90.43±4.76 | 1033.45±49.03 | 23 | ORN | 2.01±0.35 | 72.99±2.28 |
| 8 | SER | 76.83±2.81 | 1179.57±43.5 | 24 | LYS | 41.1±1.84 | 774.39±11.37 |
| 9 | ETH | 32.59±1.64 | 336.72±14.42 | 25 | α-GABA | 4.08±0.21 | 73.32±1.77 |
| 10 | GLY | 52.57±2.03 | 493.08±14.5 | 26 | CYS | 10.05±1.13 | 95.21±2.21 |
| 11 | ASP | 113.15±3.05 | 2654.44±68.02 | 27 | TYR | 96.33±2.71 | 888.9±16.47 |
| 12 | CIT | 5.72±0.39 | 75.71±0.77 | 28 | VAL | 257.05±6.12 | 2758.03±61.52 |
| 13 | SAR | 8.68±0.35 | 14.34±0.41 | 29 | ILE | 105.72±2.51 | 1829.82±47.31 |
| 14 | GLU | 233.47±6.86 | 2909.83±53.94 | 30 | LEU | 80.64±2.67 | 1600.56±28.88 |
| 15 | β-ALA | 49.73±1.22 | 368.66±5.51 | 31 | PHE | 36.64±1.25 | 603.16±13.87 |
| 16 | THR | 190.31±6.2 | 1801.33±52.7 | 32 | TRP | 68.02±3.37 | 349.4±8.68 |

As an important component of living organisms, amino acids play an important role in the regulation of material metabolism and information transfer in the body of life(Cruz et al. 2024). The composition of the amino acids in GBF was found to be balanced. Additionally, the content of γ-aminobutyric acid (GABA) in GBF was determined to be 3.6 mg/g, which was approximately 6.58 times that of GBL. Studies have demonstrated that GABA is a crucial inhibitory neurotransmitter in the central nervous system. It is an active amino acid that plays a pivotal role in the energy metabolism of the brain (Bak, Schousboe, and Waagepetersen 2006).

#### 2.2.2 Fatty Acid Content Determination

A comprehensive analysis revealed the presence of 27 fatty acids within the GBF (**Table 5**). Of these 27 fatty acids, the content of GBF was higher than that of GBL for all 26 fatty acids, except for FA28:0. The total fatty acid content of GBF was 1.95 mg/g, which was about 7.4 times that of GBL, among them Myristoleic acid (FA14:1); Myristoleic acid(FA14:1); 9-hexadecenoic acid; palmitoleic acid(FA16:1); Heptadecenoic acid(FA17:1); Oleic acid(FA18:1); Linoleic acid(FA18:2); Alpha-Linolenic acid(FA18:3); Erucic acid(FA20:1); Eicosadienoic acid(FA20:2); eicosatrienoic acid(FA20:3); Arachidonic acid(FA20:4); Eicosapentaenoic Acid(FA20:5); and Docosapentaenoic acid(FA22:5) 12 unsaturated fatty acids content of 1.333 mg/g, accounting for about 68.3% of the total fatty acid content, while the unsaturated fatty acid content in the leaf is only 0.106 mg/g, the highest content of FA18:2 in the GBF reaches 0.735 mg/g, FA18:2 can be converted to FA18:3 and FA20:4, and recently it has been shown that FA18:2 is an effective neuroprotective and anti-inflammatory agent in Parkinson’s disease models (Alarcon-Gil et al. 2022). FA18:3 (ALA) content up to 0.151 mg/g, studies have shown that ALA reduced total cholesterol, LDL cholesterol, and blood pressure, and some trials also have shown an anti-inflammatory effect of ALA, which collectively account for, in part, the cardiovascular benefits of ALA(Sala-Vila et al. 2022).

**Table 5.** Free fatty acids in GBF and GBL in various classes and contents.

| No. |  | GBL | GBF | No. |  | GBL | GBF |
| --- | --- | --- | --- | --- | --- | --- | --- |
| 1 | FA11:0 | 0.43±0.07 | 0.86±0.09 | 15 | FA 20:1 | 0.32±0.03 | 2.37±0.14 |
| 2 | FA12:0 | 2.26±0.13 | 3.77±0.24 | 16 | FA 20:2 | 1.57±0.06 | 37.4±1.38 |
| 3 | FA14:1 | 0.72±0.03 | 1.16±0.12 | 17 | FA 20:3 | 7.66±0.21 | 220.72±10.81 |
| 4 | FA14:0 | 13.47±0.43 | 15.62±1.13 | 18 | FA 20:4 | 1.1±0.01 | 10.52±0.49 |
| 5 | FA15:0 | 3.03±0.26 | 10.22±0.78 | 19 | FA 20:5 | 1.7±0.08 | 7.13±0.29 |
| 6 | FA16:0 | 109.46±3.87 | 508.51±25.08 | 20 | FA 21:0 | 0.37±0.1 | 1.35±0.17 |
| 7 | FA16:1 | 3.91±0.13 | 13.22±0.34 | 21 | FA 22:0 | 1.11±0.1 | 8.78±0.68 |
| 8 | FA17:0 | 3.41±0.03 | 16.14±0.88 | 22 | FA 22:5 | 0.71±0.14 | 1.64±0.06 |
| 9 | FA17:1 | 2.07±0.11 | 4.57±0.2 | 23 | FA 23:0 | 0.85±0.06 | 2.77±0.13 |
| 10 | FA18:0 | 11±1.26 | 21.98±2.62 | 24 | FA 24:0 | 3.29±0.08 | 13.98±0.81 |
| 11 | FA18:1 | 34.73±0.68 | 149.43±10.33 | 25 | FA 25:0 | 1.18±0 | 1.99±0.18 |
| 12 | FA18:2 | 30.71±0.65 | 734.53±26.93 | 26 | FA 26:0 | 2.94±0.14 | 4.8±0.22 |
| 13 | FA18:3 | 21.02±0.9 | 150.77±7.38 | 27 | FA 28:0 | 3.58±0.11 | 2.33±0.21 |
| 14 | FA 20:0 | 1.34±0.06 | 6.01±0.27 |  |  |  |  |

#### 2.2.3 Inorganic Element Content Analysis

In the GBF and GBL, 11 inorganic elements were detected (**Table 6**). These encompassed crucial macronutrients such as K, Ca, Mg, Fe, and P. Among the macronutrients, K; Na, and P were higher in GBF than in GBL, with the highest content of potassium at 47.43 mg/g, which was about 1.63 times that of GBL, and that of P at 7.46 mg/g, which was about 6.05 times that of GBL. Meanwhile, the content of Mg in GBL is 6.77 mg/g, which is about 2.9 times that of GBF. Mg plays a vital role in protein synthesis, energy release, and enamel strength by aiding Ca retention, and it also acts as a neurotransmitter(Arya, Salve, and Chauhan 2016). Furthermore, the GBF and GBL exhibited an assortment of trace minerals essential for human physiology, including Cu, Fe, Mn, Zn, Sr, and Al. The content of trace elements Cu, Zn, and Al in GBF is higher than GBL, with the highest content of Al being 0.028 mg/g and the content of Cu being 0.014 mg/g, which is approximately 6.76 times that of GBL.

**Table 6.** Inorganic elements in GBF and GBL in various classes and contents.

|  | GBL | GBF |
| --- | --- | --- |
| K | 29084.002 | 47438.077 |
| Na | 548.99474 | 597.98229 |
| Ca | 458.83083 | 299.83574 |
| Mg | 6768.0404 | 2333.1223 |
| Fe | 28.944048 | 28.822286 |
| Cu | 2.106819 | 14.249728 |
| Zn | 7.9813802 | 16.086908 |
| Mn | 15.98756 | 5.7050287 |
| Sr | 7.386061 | 4.3658509 |
| Al | 8.9337713 | 28.448315 |
| P | 1232.5758 | 7457.1208 |

### 2.3 Cell Activity Assay

It is established that ferroptosis occupies an important place among the forms of cell death induced by ionizing radiation. It has been found that it is possible to mitigate radiation-induced brain damage by modulating ferroptosis.

#### 2.3.1 Anti-Radiation Assays

After PC12 cells were irradiated with 8 Gy X-rays, the survival rate of PC12 cells in the IR group decreased to 66.9% compared with the Control, whereas both GBF and GBL increased the survival rate of PC12 cells to a certain extent in a dose-dependent manner. GBF at 200 µg/mL increased PC12 cell viability to 85.2%, which was superior to the positive drug Exrad (79.2%) and GBL (73.1%) (Figure 2a). The above results indicate that GBF has significant anti-radiation activity, which is superior to GBL and to some extent to the positive drug Exrad.

**Figure 2.**
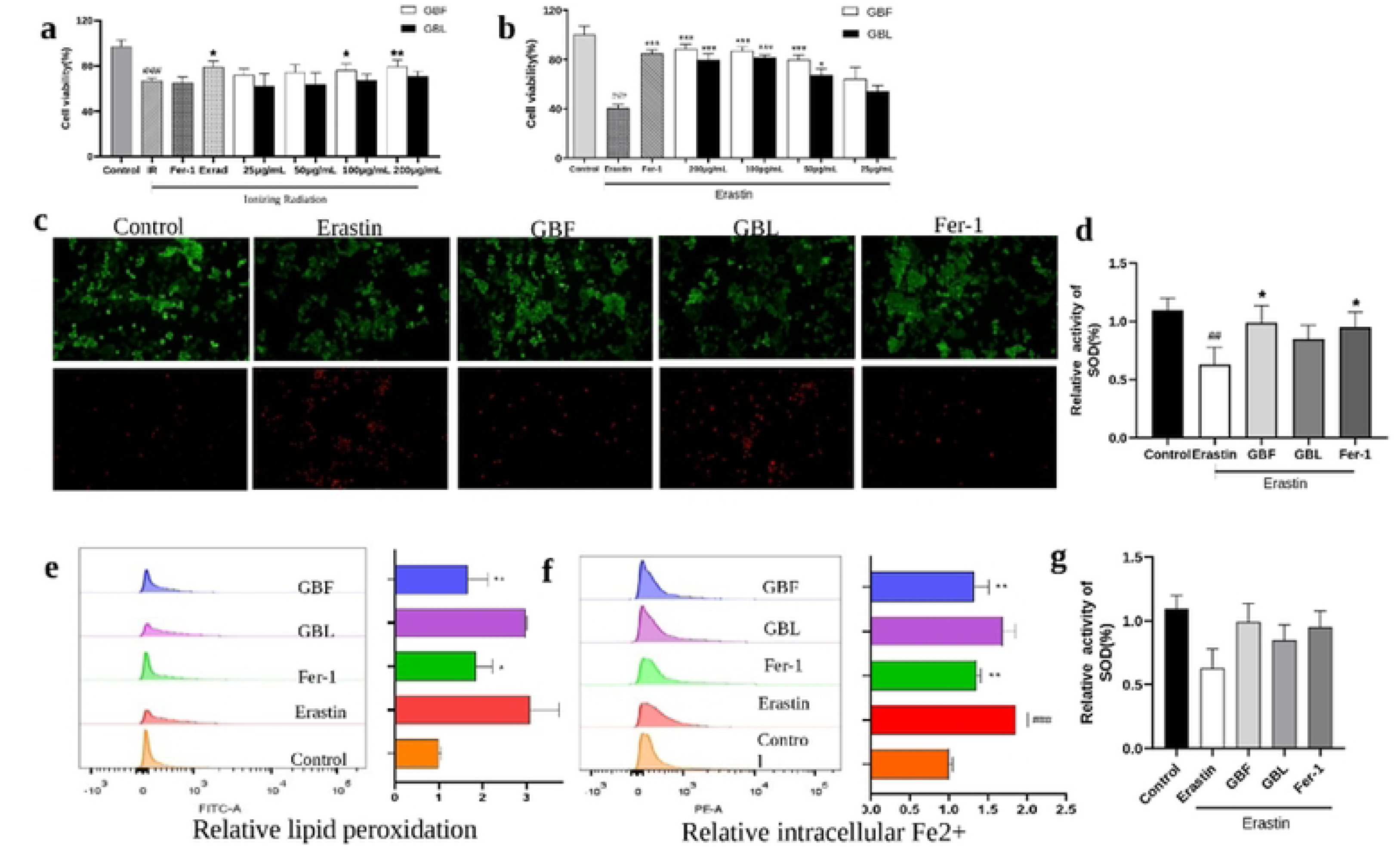
Effects of GBF and GBL on erastin-induced ferroptosis and irradiated with X-rays. Effect of GBF and GBL on the survival of PC12 irradiated with X-rays (a). Effect of GBF and GBL on the cell viability of PC12cells treated with ferroptosis inducer (b). Calcein-AM/PI staining of PC12 cells after erastin exposure. Red fluorescence indicates dead cells and green fluorescence indicates live cells (c). Flow cytometry results indicate ROS production in living PC12 cells (d). Flow cytometry results indicate lipid peroxidation in living PC12 cells (e). Flow cytometry results indicate Fe^2+^ in living PC12 cells (f). The relative activity of GSH in PC12 cells (g). (Compared with the Control group, ^###^*P*<0.001; compared with the model group \**P*<0.05, \*\*\**P*<0.001)

GBL is one of the best-selling herbal remedies in the world and the most-sold herbal supplement in the US and Europe(Biernacka, Adamska, and Felisiak 2023). However, the development of the chemical composition and pharmacological activity of the GBF is still in its infancy, so this study focuses on a comparative study of the chemical and nutritional composition and a preliminary investigation of the anti-radiation activity and mechanism of the two (Li 2019).

#### 2.3.2 Anti-Ferroptosis Assay

Erastin is a commonly used inducer of ferroptosis. Erastin reduced the viability of PC12 cells to 40.75% compared to the Control (Figure 2b). Fer-1, a well-characterized inhibitor of ferroptosis, was selected as a positive control in this study and was observed to increase cell survival to 84.8%. The addition of both GBF and GBL inhibited ferroptosis in a dose-dependent manner, and GBF increased cell survival significantly better than GBL at different concentrations, with GBF at 200 µg/mL increasing cell survival to 88.7 %. The GBL was 79.50%. To further characterize the erastin-induced cell damage, we used Calcein-AM/PI staining to distinguish between living and dead cells, noting that the ratio of dead/live cells increased gradually, GBF significantly improved this phenomenon and was superior to GBL and Fer-1 (Figure 2c).

To determine the involvement of ferroptosis in erastin-triggered PC12 cell injury, ROS as a biomarker of peroxidation of the lipids was detected with the DCFH-DA probe (Figure 2d). Additionally, we used a BODIPY™ 581/591 C11 probe to detect LPO in PC12 cells (Figure 2e), and the FeRhoNox^TM^-1 probe to detect Fe^2+^(Figure 2f). We found that ROS, LPO, and Fe^2+^ levels increased with ferroptosis occurs. Both GBF and GBL were able to reduce ROS, LPO, and Fe^2+^ levels to some extent, and GBF was more effective than GBL. Ferroptosis also caused a decrease in the levels of SOD, GBF was able to increase SOD levels, and GBF was more effective than GBL and Fer-1.

Some studies reported that IR induced ROS generation, increased iron levels, and altered expression of ferroptosis-related genes(Berry et al. 2024b). The above results indicate that GBF has the ability to significantly inhibit the ferroptosis of neuronal cells and is more effective than GBL, suggesting that GBF is able to ameliorate neuronal ferroptosis by attenuating iron-dependent lipid peroxidation in PC12 cells

## 3 Conclusion

In the present study, we found that GBF contains active components such as flavonoids and terpene lactones, chemical components such as purines and ceramides, and nutrients such as amino acids, fatty acids, and inorganic elements, which is sufficient to prove that GBF also has excellent pharmacological potential. In fact, the majority of these components are present in greater quantities than in GBL. This suggests that Ginkgo biloba flowers is a plant extract with superior benefits and enhanced potential for development into functional foods and nutraceuticals.

Ferroptosis is one of the most important forms of cell death after irradiation and GBF has good activities against radiation as well as ferroptosis, suggesting that GBF has excellent pharmacological potential and the performance of GBF was significantly better than that of GBL. GBF has good activity, which suggests that GBF probably exerts its anti-radiation effect by targeting ferroptosis.

This paper presents a preliminary differential analysis of the chemical and nutritional composition of GBF and GBL, as well as the anti-radiation activities of both, with the aim of exploring the broader activity and material basis of Ginkgo biloba for better exploitation of GBF, which is a valuable resource.

## Acknowledgements

We would like to thank Prof. Min Li and Ms. Ruihong Li for their valuable contributions to this research. Meanwhile, We would also like to thank our families for their constant support and encouragement throughout this research.

## Data availability

Data will be made available on request.

## Reference

Y. Liu, H. Xin, Y. Zhang, F. Che, N. Shen, and Y. Cui, “Leaves, seeds and exocarp of Ginkgo biloba L. (Ginkgoaceae): A Comprehensive Review of Traditional Uses, phytochemistry, pharmacology, resource utilization and toxicity,” J Ethnopharmacol, vol. 298, p. 115645, Nov. 2022, doi: 10.1016/j.jep.2022.115645.

M. Li, “Chemical constituents from the Male Flowers of Ginkgo biloba L. and Their Biological Activities,” BeiJing Institute of Radiation Medicine, BeiJing, China, 2019.

C. E. Berry et al., “Role of ferroptosis in radiation-induced soft tissue injury,” Cell Death Discov, vol. 10, p. 313, Jul. 2024, doi: 10.1038/s41420-024-02003-5.

R. Li et al., “Advances in Supercritical Carbon Dioxide Extraction of Bioactive Substances from Different Parts of Ginkgo biloba L.,” Molecules, vol. 26, no. 13, p. 4011, Jun. 2021, doi: 10.3390/molecules26134011.

null Noor-E-Tabassum, et al., “Ginkgo biloba: A Treasure of Functional Phytochemicals with Multimedicinal Applications,” Evid Based Complement Alternat Med, vol. 2022, p. 8288818, 2022, doi: 10.1155/2022/8288818.

L. E. B. Bettio, J. Gil-Mohapel, and A. L. S. Rodrigues, “Guanosine and its role in neuropathologies,” Purinergic Signal, vol. 12, no. 3, pp. 411–426, Sep. 2016, doi: 10.1007/s11302-016-9509-4.

A. Schwarz and A. H. Futerman, “Distinct Roles for Ceramide and Glucosylceramide at Different Stages of Neuronal Growth,” J Neurosci, vol. 17, no. 9, pp. 2929–2938, May 1997, doi: 10.1523/JNEUROSCI.17-09-02929.1997.

T. M. Cruz et al., “Bioaccessibility of bioactive compounds from Pereskia aculeata and their cellular antioxidant effect,” Food Chem, vol. 460, no. Pt 1, p. 140484, Dec. 2024, doi: 10.1016/j.foodchem.2024.140484.

L. K. Bak, A. Schousboe, and H. S. Waagepetersen, “The glutamate/GABA-glutamine cycle: aspects of transport, neurotransmitter homeostasis and ammonia transfer,” Journal of Neurochemistry, vol. 98, no. 3, pp. 641–653, Aug. 2006, doi: 10.1111/j.1471-4159.2006.03913.x.

J. Alarcon-Gil et al., “Neuroprotective and Anti-Inflammatory Effects of Linoleic Acid in Models of Parkinson’s Disease: The Implication of Lipid Droplets and Lipophagy,” Cells, vol. 11, no. 15, p. 2297, Jul. 2022, doi: 10.3390/cells11152297.

A. Sala-Vila, J. Fleming, P. Kris-Etherton, and E. Ros, “Impact of α-Linolenic Acid, the Vegetable ω-3 Fatty Acid, on Cardiovascular Disease and Cognition,” Adv Nutr, vol. 13, no. 5, pp. 1584–1602, Oct. 2022, doi: 10.1093/advances/nmac016.

S. S. Arya, A. R. Salve, and S. Chauhan, “Peanuts as functional food: a review,” J Food Sci Technol, vol. 53, no. 1, pp. 31–41, Jan. 2016, doi: 10.1007/s13197-015-2007-9.

P. Biernacka, I. Adamska, and K. Felisiak, “The Potential of Ginkgo biloba as a Source of Biologically Active Compounds—A Review of the Recent Literature and Patents,” Molecules, vol. 28, no. 10, p. 3993, May 2023, doi: 10.3390/molecules28103993.

C. E. Berry et al., “Role of ferroptosis in radiation-induced soft tissue injury,” Cell Death Discovery, vol. 10, p. 313, Jul. 2024, doi: 10.1038/s41420-024-02003-5.

Y. Zhang, S. Zhu, Y. Gu, Y. Feng, and B. Gao, “Network Pharmacology Combined with Experimental Validation to Investigate the Mechanism of the Anti-Hyperuricemia Action of Portulaca oleracea Extract,” Nutrients, vol. 16, no. 20, p. 3549, Oct. 2024, doi: 10.3390/nu16203549.

ShangGuan X., “Application of spectrophotometry in drug analysis - A study on the analytical methods of ginkgo biloba flavonoids and nicotinamide,” Master, Northwest University, 2004. Accessed: Nov. 01, 2024. [Online]. Available: https://kns.cnki.net/KCMS/detail/detail.aspx?dbcode=CMFD&dbname=CMFD9904&filename=2004104450.nh

Y. Sawada et al., “Widely Targeted Metabolomics Based on Large-Scale MS/MS Data for Elucidating Metabolite Accumulation Patterns in Plants,” Plant and Cell Physiology, vol. 50, no. 1, p. 37, Dec. 2008, doi: 10.1093/pcp/pcn183.

S. Repert, S. Matthes, and W. Rozhon, “Quantification of Arbutin in Cosmetics, Drugs and Food Supplements by Hydrophilic-Interaction Chromatography,” Molecules, vol. 27, no. 17, p. 5673, Sep. 2022, doi: 10.3390/molecules27175673.

M. C. Roman, J. M. Betz, and J. Hildreth, “Determination of Synephrine in Bitter Orange Raw Materials, Extracts, and Dietary Supplements by Liquid Chromatography with Ultraviolet Detection: Single-Laboratory Validation,” Journal of AOAC International, vol. 90, no. 1, p. 68, Feb. 2007.

W. K, I. S, U. K, S. M, and H. M, “An antivitamin B6, 4’-methoxypyridoxine, from the seed of Ginkgo biloba L.,” Chemical & Pharmaceutical Bulletin, vol. 33, no. 8, pp. 3555–3557, Jan. 1985, doi: 10.1248/CPB.33.3555.

R. Kumar, G. Murugananthan, K. Nandakumar, and S. Talwar, “Isolation of anxiolytic principle from ethanolic root extract of Cardiospermum halicacabum,” Phytomedicine, vol. 18, no. 2–3, pp. 219–223, Jan. 2011, doi: 10.1016/j.phymed.2010.07.002.

J. Yu et al., “Non-target metabolomics unravels the effect and mechanism of Lianpu Drink on spleen-stomach damp-heat syndrome,” J Chromatogr B Analyt Technol Biomed Life Sci, vol. 1246, p. 124281, Oct. 2024, doi: 10.1016/j.jchromb.2024.124281.

J. Cao, H. Wang, W. Zhang, F. Cao, G. Ma, and E. Su, “Tailor-Made Deep Eutectic Solvents for Simultaneous Extraction of Five Aromatic Acids from Ginkgo biloba Leaves,” Molecules, vol. 23, no. 12, p. 3214, Dec. 2018, doi: 10.3390/molecules23123214.

E. Mfotie Njoya, L. J. McGaw, and T. J. Makhafola, “Investigating the Phytochemical Composition, Antioxidant, and Anti-Inflammatory Potentials of Cassinopsis ilicifolia (Hochst.) Kuntze Extract against Some Oxidative Stress and Inflammation Molecular Markers,” Curr Issues Mol Biol, vol. 46, no. 9, pp. 9639–9658, Sep. 2024, doi: 10.3390/cimb46090573.

K. Tawaha, R. Sadi, F. Qa’dan, K. Z. Matalka, and A. Nahrstedt, “A bioactive prodelphinidin from Mangifera indica leaf extract,” Z Naturforsch C J Biosci, vol. 65, no. 5–6, pp. 322–326, 2010, doi: 10.1515/znc-2010-5-603.

M. Dou et al., “HPLC combined with chemometrics for quality control of Huamoyan Granules or Capsules,” Chin Herb Med, vol. 16, no. 3, pp. 449–456, Jul. 2024, doi: 10.1016/j.chmed.2023.03.005.

K. Js et al., “Constituents from leaves of Macaranga hemsleyana,” Chinese herbal medicines, vol. 16, no. 3, Aug. 2023, doi: 10.1016/j.chmed.2023.03.006.

Y. He et al., “Quality Markers’ Discovery and Quality Evaluation of Jigucao Capsule Using UPLC-MS/MS Method,” Molecules, vol. 28, no. 6, p. 2494, Mar. 2023, doi: 10.3390/molecules28062494.

PubChem, “3,4,4a,5,6,7-Hexahydro-1,1,4a-trimethyl-2(1H)-naphthalenone.” Accessed: Nov. 05, 2024. [Online]. Available: https://pubchem.ncbi.nlm.nih.gov/substance/275191875

X. Chen, L. Kong, L. Sheng, X. Li, and H. Zou, “[Applications of two-dimensional liquid chromatography coupled to mass spectrometry for the separation and identification of compounds in ginkgo biloba extracts],” Se Pu, vol. 23, no. 1, pp. 46–51, Jan. 2005.

F. Xu et al., “Isolation, characterization, and function analysis of a flavonol synthase gene from Ginkgo biloba,” Mol Biol Rep, vol. 39, no. 3, pp. 2285–2296, Mar. 2012, doi: 10.1007/s11033-011-0978-9.

P. Biernacka, K. Felisiak, and I. Adamska, “The potential of dried Ginkgo Biloba leaves as a novel ingredient in fermented beverages of enhanced flavour and antioxidant properties,” Food Chem, vol. 461, p. 141018, Dec. 2024, doi: 10.1016/j.foodchem.2024.141018.

S. Gao et al., “Evaluation of the Effects of Processing Technique on Chemical Components in Raphani Semen by HPLC and UPLC-Q-TOF-MS,” Int J Anal Chem, vol. 2022, p. 8279839, 2022, doi: 10.1155/2022/8279839.

M. Ellnain-Wojtaszek and G. Zgórka, “HIGH-PERFORMANCE LIQUID CHROMATOGRAPHY AND THIN-LAYER CHROMATOGRAPHY OF PHENOLIC ACIDS FROM GINKGO BILOBA L. LEAVES COLLECTED WITHIN VEGETATIVE PERIOD,” Journal of Liquid Chromatography & Related Technologies, Jan. 1999, doi: 10.1081/JLC-100101744.

J. Wu et al., “A Flavonoid Glycoside Compound from Siraitia grosvenorii with Anti-Inflammatory and Hepatoprotective Effects In Vitro,” Biomolecules, vol. 14, no. 4, p. 450, Apr. 2024, doi: 10.3390/biom14040450.

H. Wang, L. Zeng, and H. Ding, “A Novel Strategy for Screening of Secondary Metabolites in Ginkgo biloba Leaves by Ultra-Performance Liquid Chromatography–Tandem Information Dependent Acquisition Mass Spectrometry,” J Anal Chem, vol. 79, no. 3, pp. 330–338, Mar. 2024, doi: 10.1134/S1061934824030146.

T. R. Baker and B. T. Regg, “A multi-detector chromatographic approach for characterization and quantitation of botanical constituents to enable in silico safety assessments,” Analytical and Bioanalytical Chemistry, vol. 410, no. 21, p. 5143, Jul. 2018, doi: 10.1007/s00216-018-1163-y.

M. Meenu et al., “Impact of inherent chemical composition of wheat and various processing technologies on whole wheat flour and its final products,” CEREAL RESEARCH COMMUNICATIONS, Jun. 2024, doi: 10.1007/s42976-024-00544-0.

M. Shirai, Y. Kawai, R. Yamanishi, and J. Terao, “Approach to novel functional foods for stress control 5. Antioxidant activity profiles of antidepressant herbs and their active components,” J Med Invest, vol. 52 Suppl, pp. 249–251, Nov. 2005, doi: 10.2152/jmi.52.249.

F. Zhang et al., “LC-MS based strategy for chemical profiling and quantification of dispensing granules of Ginkgo biloba seeds,” Heliyon, vol. 10, no. 17, p. e36909, Sep. 2024, doi: 10.1016/j.heliyon.2024.e36909.

Y. Sun et al., “Investigation on the mechanism of Ginkgo Folium in the treatment of Non-alcoholic Fatty Liver Disease by strategy of network pharmacology and molecular docking,” Technol Health Care, vol. 31, no. S1, pp. 209–221, 2023, doi: 10.3233/THC-236018.

D. Lee et al., “Identification of Anti-Inflammatory Compounds from Hawaiian Noni (Morinda citrifolia L.) Fruit Juice,” Molecules, vol. 25, no. 21, p. 4968, Oct. 2020, doi: 10.3390/molecules25214968.

K. Zhou, S. Xiao, S. Cao, C. Zhao, M. Zhang, and Y. Fu, “Improvement of glucolipid metabolism and oxidative stress via modulating PI3K/Akt pathway in insulin resistance HepG2 cells by chickpea flavonoids,” Food Chem X, vol. 23, p. 101630, Oct. 2024, doi: 10.1016/j.fochx.2024.101630.

M. I. Alruwad et al., “Insights into Clematis cirrhosa L. Ethanol Extract: Cytotoxic Effects, LC-ESI-QTOF-MS/MS Chemical Profiling, Molecular Docking, and Acute Toxicity Study,” Pharmaceuticals (Basel), vol. 17, no. 10, p. 1347, Oct. 2024, doi: 10.3390/ph17101347.

E. Bampali, S. Germer, R. Bauer, and Ž. Kulić, “HPLC-UV/HRMS methods for the unambiguous detection of adulterations of Ginkgo biloba leaves with Sophora japonica fruits on an extract level,” Pharmaceutical Biology, vol. 59, no. 1, p. 436, Apr. 2021, doi: 10.1080/13880209.2021.1910717.

H. Jiao et al., “Metabolomics Analysis of Phenolic Composition and Content in Five Pear Cultivars Leaves,” Plants (Basel), vol. 13, no. 17, p. 2513, Sep. 2024, doi: 10.3390/plants13172513.

V. K. Phan et al., “Antioxidant activity of a new C-glycosylflavone from the leaves of Ficus microcarpa,” Bioorg Med Chem Lett, vol. 21, no. 2, pp. 633–637, Jan. 2011, doi: 10.1016/j.bmcl.2010.12.025.

J. Penna-Coutinho, A. C. Aguiar, and A. U. Krettli, “Commercial drugs containing flavonoids are active in mice with malaria and in vitro against chloroquine-resistant Plasmodium falciparum,” Mem Inst Oswaldo Cruz, vol. 113, no. 12, p. e180279, Dec. 2018, doi: 10.1590/0074-02760180279.

A. Hasler, G. A. Gross, B. Meier, and O. Sticher, “Complex flavonol glycosides from the leaves of Ginkgo biloba,” Phytochemistry, vol. 31, no. 4, pp. 1391–1394, Apr. 1992, doi: 10.1016/0031-9422(92)80298-s.

B. Ma et al., “Combining metabolomics and transcriptomics to reveal the potential medicinal value of rare species Glycyrrhiza squamulose,” Heliyon, vol. 10, no. 10, p. e30868, May 2024, doi: 10.1016/j.heliyon.2024.e30868.

X. Zhang et al., “Comparative study on chemical constituents of different medicinal parts of Lonicera japonica Thunb. Based on LC-MS combined with multivariate statistical analysis,” Heliyon, vol. 10, no. 12, p. e31722, Jun. 2024, doi: 10.1016/j.heliyon.2024.e31722.

D. Aoki et al., “Distribution of coniferin in freeze-fixed stem of Ginkgo biloba L. by cryo-TOF-SIMS/SEM,” Sci Rep, vol. 6, p. 31525, Aug. 2016, doi: 10.1038/srep31525.

H. Kimura, H. Irie, K. Ueda, and S. Ueo, “[The constituents of the heartwood of Ginkgo biloba L. V. The structure and absolute configuration of bilobanone],” Journal of the Pharmaceutical Society of Japan, vol. 88, no. 5, pp. 562–572, May 1968, doi: 10.1248/YAKUSHI1947.88.5_562.

T. Zhu, L. Wang, Y. Feng, G. Sun, and X. Sun, “Classical Active Ingredients and Extracts of Chinese Herbal Medicines: Pharmacokinetics, Pharmacodynamics, and Molecular Mechanisms for Ischemic Stroke,” Oxid Med Cell Longev, vol. 2021, p. 8868941, 2021, doi: 10.1155/2021/8868941.

H. Otsuka, X. N. Zhong, E. Hirata, T. Shinzato, and Y. Takeda, “Myrsinionosides A-E: megastigmane glycosides from the leaves of Myrsine seguinii Lev,” Chem Pharm Bull (Tokyo), vol. 49, no. 9, pp. 1093–1097, Sep. 2001, doi: 10.1248/cpb.49.1093.

J. A. Kim et al., “New monoterpene glycosides and phenolic compounds from Distylium racemosum and their inhibitory activity against ribonuclease H,” Bioorg Med Chem Lett, vol. 21, no. 10, pp. 2840–2844, May 2011, doi: 10.1016/j.bmcl.2011.03.091.

P. Songvut et al., “A validated LC-MS/MS method for clinical pharmacokinetics and presumptive phase II metabolic pathways following oral administration of Andrographis paniculata extract,” Sci Rep, vol. 13, no. 1, p. 2534, Feb. 2023, doi: 10.1038/s41598-023-28612-1.

M. M. Lephatsi, M. S. Choene, A. P. Kappo, N. E. Madala, and F. Tugizimana, “An Integrated Molecular Networking and Docking Approach to Characterize the Metabolome of Helichrysum splendidum and Its Pharmaceutical Potentials,” Metabolites, vol. 13, no. 10, p. 1104, Oct. 2023, doi: 10.3390/metabo13101104.

T. Yang, F. Deng, X. Yang, and S. Li, “Establishment of HPLC Fingerprints for Feiqizhong Tablets and Simultaneous Determination of Fourteen Constituents,” J Anal Methods Chem, vol. 2024, p. 7703951, 2024, doi: 10.1155/2024/7703951.

Y. Wang, M. Jia, Y. Gao, and B. Zhao, “Multiplex Quantitative Analysis of 9 Compounds of Scutellaria baicalensis Georgi in the Plasma of Respiratory Syncytial Virus-Infected Mice Based on HPLC-MS/MS and Pharmacodynamic Effect Correlation Analysis,” Molecules, vol. 28, no. 16, p. 6001, Aug. 2023, doi: 10.3390/molecules28166001.

J. Qiu et al., “Screening and Identifying Antioxidative Components in Ginkgo biloba Pollen by DPPH-HPLC-PAD Coupled with HPLC-ESI-MS2,” PLoS ONE, vol. 12, no. 1, p. e0170141, Jan. 2017, doi: 10.1371/journal.pone.0170141.

R. N. Muchiri and R. B. van Breemen, “Chemical Standardization of Milk Thistle (Silybum marianum L.) Extract Using UHPLC-MS/MS and the Method of Standard Addition,” J Am Soc Mass Spectrom, vol. 35, no. 8, pp. 1726–1732, Aug. 2024, doi: 10.1021/jasms.4c00125.

J. Ni et al., “Transgenic tobacco plant overexpressing ginkgo dihydroflavonol 4-reductase gene GbDFR6 exhibits multiple developmental defects,” Front Plant Sci, vol. 13, p. 1066736, 2022, doi: 10.3389/fpls.2022.1066736.

R. Mao et al., “Transcriptome and HPLC Analysis Reveal the Regulatory Mechanisms of Aurantio-Obtusin in Space Environment-Induced Senna obtusifolia Lines,” Int J Environ Res Public Health, vol. 19, no. 2, p. 898, Jan. 2022, doi: 10.3390/ijerph19020898.

S. K. Hyun, S. S. Kang, K. H. Son, H. Y. Chung, and J. S. Choi, “Biflavone glucosides from Ginkgo biloba yellow leaves,” Chem Pharm Bull (Tokyo), vol. 53, no. 9, pp. 1200–1201, Sep. 2005, doi: 10.1248/cpb.53.1200.

Q. Zhang et al., “Application of GC/MS-based metabonomic profiling in studying the lipid-regulating effects of Ginkgo biloba extract on diet-induced hyperlipidemia in rats,” Acta Pharmacol Sin, vol. 30, no. 12, pp. 1674–1687, Dec. 2009, doi: 10.1038/aps.2009.173.

S. S. Qazi, D. A. Lombardo, and M. M. Abou-Zaid, “A Metabolomic and HPLC-MS/MS Analysis of the Foliar Phenolics, Flavonoids and Coumarins of the Fraxinus Species Resistant and Susceptible to Emerald Ash Borer,” Molecules, vol. 23, no. 11, p. 2734, Oct. 2018, doi: 10.3390/molecules23112734.

N. Bouzenad et al., “Exploring Bioactive Components and Assessing Antioxidant and Antibacterial Activities in Five Seaweed Extracts from the Northeastern Coast of Algeria,” Mar Drugs, vol. 22, no. 6, p. 273, Jun. 2024, doi: 10.3390/md22060273.

Q.-L. Chen, Z.-Y. Shi, Y.-H. Zhang, and J.-B. Zheng, “[Study on the chemical constituents in roots of Gentiana dahurica],” Zhong Yao Cai, vol. 34, no. 8, pp. 1214–1216, Aug. 2011.

O. J. A. Alves et al., “HPLC method for quantifying verbascoside in Stizophyllum perforatum and assessment of verbascoside acute toxicity and antileishmanial activity,” Front Plant Sci, vol. 14, p. 1324680, 2023, doi: 10.3389/fpls.2023.1324680.

Y. Huang et al., “Identification of key phenolic compounds for alleviating gouty inflammation in edible chrysanthemums based on spectrum-effect relationship analyses,” Food Chem X, vol. 20, p. 100897, Dec. 2023, doi: 10.1016/j.fochx.2023.100897.

X. Xu, X. Li, S. Chen, Y. Liang, C. Zhang, and Y. Huang, “Simultaneous Qualitative and Quantitative Analyses of 41 Constituents in Uvaria macrophylla Leaves Screen Antioxidant Quality-Markers Using Database-Affinity Ultra-High-Performance Liquid Chromatography with Quadrupole Orbitrap Tandem Mass Spectrometry,” Molecules, vol. 29, no. 20, p. 4886, Oct. 2024, doi: 10.3390/molecules29204886.

J. Alqahtani et al., “Potential Surviving Effect of Cleome droserifolia Extract against Systemic Staphylococcus aureus Infection: Investigation of the Chemical Content of the Plant,” Antibiotics (Basel), vol. 13, no. 5, p. 450, May 2024, doi: 10.3390/antibiotics13050450.

J. Wei, R. Xu, Y. Zhang, L. Zhao, S. Li, and Z. Zhao, “Ultra-High-Performance Liquid Chromatography-Electrospray Ionization-High-Resolution Mass Spectrometry for Distinguishing the Origin of Ellagic Acid Extracts: Pomegranate Peels or Gallnuts,” Molecules, vol. 29, no. 3, p. 666, Jan. 2024, doi: 10.3390/molecules29030666.

W. Siheri et al., “Isolation of a Novel Flavanonol and an Alkylresorcinol with Highly Potent Anti-Trypanosomal Activity from Libyan propolis,” Molecules, vol. 24, no. 6, p. 1041, Mar. 2019, doi: 10.3390/molecules24061041.

F. Oiram Filho, M. P. Mitri, G. J. Zocolo, K. M. Canuto, and E. S. de Brito, “Validation of a Method for Anacardic Acid Quantification in Cashew Peduncles via High-Performance Liquid Chromatography Coupled to a Diode-Array Detector,” Foods, vol. 12, no. 14, p. 2759, Jul. 2023, doi: 10.3390/foods12142759.

J. Irie, M. Murata, and S. Homma, “Glycerol-3-phosphate Dehydrogenase Inhibitors, Anacardic Acids, from Ginkgo biloba,” Biosci Biotechnol Biochem, vol. 60, no. 2, pp. 240–243, Jan. 1996, doi: 10.1271/bbb.60.240.

